# Quantification of all 12 canonical ribonucleotides by real-time fluorogenic *in vitro* transcription

**DOI:** 10.1101/2023.02.18.527797

**Authors:** Janne Purhonen, Jukka Kallijärvi

## Abstract

Enzymatic methods to quantify deoxyribonucleoside triphosphates have existed for decades. In contrast, no general enzymatic method to quantify ribonucleoside triphosphates (rNTPs), which drive almost all cellular processes and serve as precursors of RNA, exists to date. ATP can be measured with an enzymatic luminometric method employing the firefly luciferase, but the quantification of the other ribonucleoside mono-, di-, and triphosphates is still a challenge for a non-specialized laboratory and practically impossible without chromatography equipment. To allow feasible quantification of ribonucleoside phosphates in any laboratory with typical molecular biology and biochemistry tools, we developed a robust microplate assay based on real-time detection of the Broccoli aptamer during *in vitro* transcription. The assay employs the bacteriophage T7 and SP6 RNA polymerases, two oligonucleotide templates encoding the 49-nucleotide Broccoli aptamer, and a high-affinity fluorogenic aptamer-binding dye to quantify each of the four canonical rNTPs. The inclusion of nucleoside mono- and diphosphate kinases in the assay reactions enabled the quantification of the mono- and diphosphate counterparts. The assay is inherently specific and tolerates concentrated tissue and cell extracts. In summary, we describe the first chromatography-free method to quantify ATP, ADP, AMP, GTP, GDP, GMP, UTP, UDP, UMP, CTP, CDP, and CMP in biological samples.

## INTRODUCTION

Nucleotides form genetic information, drive almost all cellular processes, and are also essential signalling molecules and biosynthetic precursors (1). Information on their cellular concentrations and phosphorylation statuses is increasingly needed in various fields of life sciences. Chromatography-based methods for the quantification of nucleotides have existed for decades (2). In addition, DNA polymerase-based methods for the determination of deoxyribonucleoside triphosphates (dNTPs) have been available since 1969 (3). Nevertheless, reliable nucleotide quantification from complex biological samples is still a challenge, especially for a non-specialized laboratory (4). Recently, we published an improved highly sensitive DNA polymerase-based method for convenient quantification of dNTPs in high-throughput manner in 384-well microplates (5). This method allows dNTP quantification in any laboratory with a typical qPCR instrument. dNTPs are, however, a minor cellular nucleotide pool needed mainly for DNA repair and replication, and of interest mainly to specialized research niches (6). Ribonucleoside triphosphates (rNTPs) are a several orders of magnitude more abundant class of cellular nucleotides than dNTPs. ATP, the most abundant of them, is easily quantified using a luminometric method employing the firefly luciferase (7). A similarly convenient method for the quantification of the other three canonical rNTPs (GTP, UTP, and CTP) and the corresponding nucleoside mono- and diphosphates (rNMPs and rNDPs, respectively) is lacking.

Analogous to the DNA polymerase-based methods for dNTPs (5), a partly similar approach could in principle allow the quantification of rNTPs with RNA polymerases. However, to the best of our knowledge, such a method has never been reported. One obstacle in the development of RNA polymerase-based method may have been the lack of practical quantitative readout. Unlike DNA polymerases, RNA polymerases undergo cycles of abortive initiation, leading to short (<14 nt) truncated RNA products (8). These short RNA products surpass the number of full-length transcripts when one rNTP is available at a rate-limiting concentration (9). Thus, the total RNA produced or labelled non-limiting nucleotides incorporated are unlikely to serve as sensitive readouts. Since the identification of several fluorogenic dye-binding RNA aptamers (RNA sequences naturally folding into a defined conformation), quantitative *in vitro* transcription with real-time detection has now, however, become possible (10, 11). Here, we utilized the highly sensitive fluorometric detection of the Broccoli aptamer (12) for the development of an RNA polymerase-based method for the quantification of rNTPs and the corresponding mono- and diphosphates.

## MATERIALS AND METHODS

### Oligonucleotides and other reagents

Integrated DNA Technologies (IDT) (Coralville, Iowa, USA) synthetized all DNA oligonucleotides used in this study (Table 1). The oligonucleotides longer than 50 nt were synthetised using IDT Ultramer technology. Standard desalting was applied to all synthetic oligonucleotides. Double-stranded DNA templates were prepared by heating equimolar mix of the complementary DNA strands to 95°C and by slowly cooling the mixture to 23-30°C.

**Table 1.**
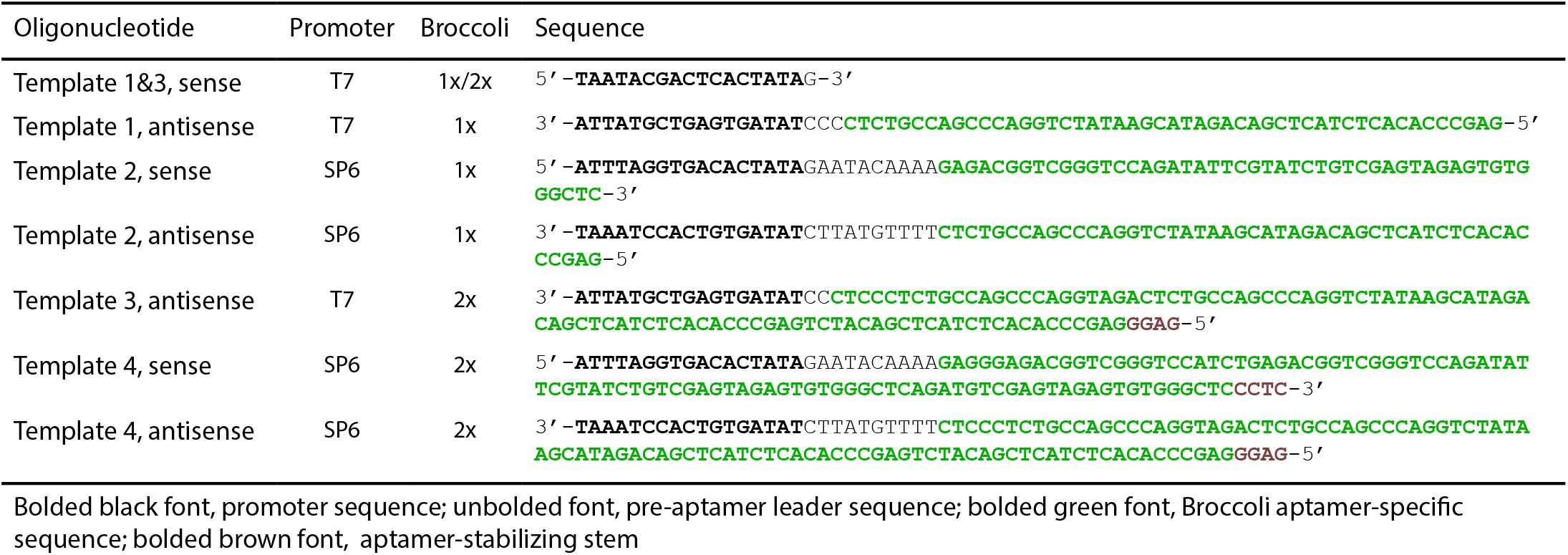
DNA templates.

Supplementary Table S2 lists all essential reagents. The following molar extinction coefficients (mM^-1^cm^-1^) were used to determine the concentrations of the nucleotide standards: 15.4 (259 nm) adenosine, 10 (262 nm) uridine, 8.9 (271 nm) cytidine, and 13.7 (252 nm) guanosine.

### Mouse tissues samples

Mice of congenic C57BL/6JCrl background were maintained by Laboratory Animal Center of the University of Helsinki. Two-month-old male mice were euthanized by cervical dislocation, and the liver tissue was immediately (10-20s) excised and placed in liquid nitrogen and stored at 80°C. For experiments with delayed sample collection, the excised liver was kept at room temperature for the indicated periods.

### Cell culture

The mouse hepatocyte line AML12 was maintained in Dulbecco’s modified Eagle’s medium-F12 mixture (DMEM-F12) with 5 mM glucose and supplemented with 10% fetal bovine serum, penicillin and streptomycin, 2 mM L-alanyl-L-glutamine, 10 μg/ml insulin, 5.5 μg/ml transferrin, and 5 ng/ml selenium. Mouse hepatoma Hepa1-6 cells were maintained in DMEM with 25 mM glucose, 10% fetal bovine serum, penicillin and streptomycin, and 2 mM L-alanyl-L-glutamine. Both cell lines were cultured at 37°C in an atmosphere of 5% CO_2_ and 95% air. For treatments, the cells were plated in 6-cm plates and grown to 60-80% confluence. The experimental chemicals were added directly to the medium and the cells were incubated for 6 hours. To collect the samples, the cells were rinsed once with cold PBS, 0.5 ml of ice-cold 80 % MeOH was added, and the cells were scraped off the plates. The cell suspensions were stored at -80°C until the extraction.

### Extraction of nucleotides

Supplementary methods describe an extraction protocol that was employed for the generation of data in Figure 4A and Supplementary Figures S3 and S4. Here, an optimized extraction with a more stringent deproteination is described. Frozen liver samples (18-22 mg) were directly homogenized in 0.5 ml of ice-cold 80% MeOH with a battery-operated microtube pestle for 10s. The homogenization was finalized with a microtip probe sonicator (12s, amplitude 12%, Branson Digital Sonifier 250). Cultured cells scraped into 80% MeOH were directly sonicated. The samples were kept on dry ice between the homogenization steps. The methanolic extracts were incubated for 3 min at 100°C. Per 0.5 ml of 80% MeOH, 200 µl of chloroform was added to further precipitate macromolecules and to remove non-polar metabolites. Phase separation was induced by addition of 200 µl water. The extracts were vortexed for 10s followed by vigorous shaking for 30s. After centrifugation (3 min 180000g at 0°C), the aqueous phase was transferred into a new microtube. MeOH was removed by washing three times with 1.2 ml of diethyl ether. Any remaining layer of diethyl ether was evaporated by 10s flow of N_2_ gas. The traces of the diethyl ether from the aqueous phase were evaporated in Genevac miVac Duo centrifugal vacuum evaporator for 12 min. The remaining liquid volume after evaporation was estimated by weighing the extracts and assuming the density of 1 mg/µl. Instead of diethyl ether washes, the cell extracts were evaporated to dryness. The extracts were stored at -80°C.

### Extraction of total protein fraction

Precipitated proteins from the nucleotide extraction were sedimented by addition of 800 µl MeOH and centrifugation (3 min 18000g at 4°C). The protein pellets were dissolved in Laemmli buffer (2% SDS, 1% β-mercaptoethanol, 60 mM Tris-Cl pH 6.8, and 12% glycerol) by sonication and 5 min incubation at 95°C. A modified Bradford reagent containing 2.5 mg/ml α-cyclodextrin to chelate the interfering SDS was employed to measure protein concentrations (13). Bovine serum albumin was used as a reference protein.

### Quantification of rNTPs by *in vitro* transcription

The assay parameters were varied and optimized throughout this study. The final established assay conditions are described here. T7 RNA polymerase (#M0251, New England BioLabs) and template 3 were the most suitable combination for the quantification of ATP, UTP and CTP. SP6 RNA polymerase (#M0207, New England BioLabs) and template 4 were more optimal for the quantification of GTP. The optimized assay reactions comprised 1x CutSmart buffer (New England BioLabs) (20 mM Tris-acetate pH 7.9, 50 mM potassium acetate, 10 mM Mg-acetate, and 0.1 mg/ml BSA), 2 mM spermidine, 1 mM of each non-limiting rNTP, 1-2 µM limiting rNTP, 5 mM DTT, 20 nM DNA template, 20 µM BI, 0.5 kU/ml RiboLock (EO0382, Thermo Scientific), 0.25 U/ml pyrophosphatase (EF0221, Thermo Scientific), and 1.9 kU/ml T7 RNA polymerase or 0.5 kU/ml SP6 RNA polymerase. BI stands for Broccoli aptamer ligand (5Z)-3-(1H-Benzimidazol-7-ylmethyl)-5-[(3,5-di-fluoro-4-hydroxyphenyl)methylene]-3,5-dihydro-2-methyl-4H-imidazol-4-one dihydrochloride) (Cat. No. 7466, Tocris) (14). Some experiments were performed without the ribonuclease inhibitor (Ribolock) as this reagent was found dispensable when adequate measures to prevent ribonuclease contamination were followed. The optimal basal concentration of limiting rNTP was 1 µM for the quantification of ATP, UTP, and CTP, and 2 µM for GTP. A master mix of the reagents was prepared and mixed with samples in 384-well qPCR plate (#HSP3805, BioRad). The reactions were carried out in volumes of 4 to 10 µl. In our final established protocol, the reactions comprised one part of sample and one part of 2x master mix. The reaction set-up was prepared on ice, the plate sealed and briefly centrifuged. The Broccoli aptamer-BI fluorescence was measured upon 470 nm excitation and 510 nm emission in Synergy H1 plate reader (Biotek) for 2 to 8h at 37°C. When accurate temperature control was essential, the plates were read in Bio-Rad CFX384 qPCR instrument instead. Baseline-subtracted end-point fluorescence values were used to generate the standard curves.

### Quantification of rNMPs and rNDPs

The sum of rNDPs + rNTPs of interest were quantified by inclusion of 5 µg/ml of recombinant human nucleoside diphosphate kinase (#NDPK-33, BioNukleo GmbH, Berlin, Germany) in the assay reactions. The sum of target rNMPs + rNDPs + rNTPs were quantified by including the nucleoside diphosphate kinase and 5 µg/ml nucleoside monophosphate kinase in the assay reactions. The following recombinant nucleoside monophosphate kinases produced by BioNukleo were used: AMP kinase (#NMPK-01), GMP kinase (#NMPK-21), and UMP-CMP kinase (#NMPK-22). The concentrations of specific nucleotide species were deduced from three measurements: 1) rNTP, 2) rNTP + rNDP, and 3) rNTP+rNDP+rNMP.

### Luciferin-luciferase assay for ATP

Invitrogen ATP Determination Kit (#A22066) was employed to measure ATP by the firefly luciferase method.

### Protocol details

Step-by-step protocols are provided as supplementary files.

## RESULTS

### Development of *in vitro* transcription assay for rNTP quantification

As a starting point we tested an *in vitro* transcription protocol with fluorescent real-time monitoring by *Kartje et al*. (10). This assay utilizes T7 RNA polymerase and Broccoli aptamer and its fluorogenic ligand DFHBI-1T (3,5-difluoro-4hydroxy-benzylidene imidazolinone). While this system functions well for its purpose, characterization and optimization of *in vitro* transcription for maximum RNA yield, it lacked sensitivity required for the robust quantification of rNTPs (Figures 1A, B, blue lines). To improve the sensitivity, we substituted DFHBI-1T with BI as the Broccoli ligand. BI has a higher affinity for the aptamer and enhanced fluorogenicity in comparison to DFHBI-1T, allowing imaging of even single Broccoli-tagged transcripts in live cells (14). It also promotes the folding and thermal stability of the aptamer. The reaction buffer of Kartje et al. assay includes 5 mM NaCl and 1 mM KCl as enhancers of aptamer folding (10), while the efficient folding of Broccoli aptamers requires ∼50 mM K^+^ or more than 200 mM Na^+^ (15). Therefore, we chose a buffer containing 50 mM potassium acetate. These changes notably increased the aptamer fluorescence (Figures 1A and B). In these initial test runs, we used a rather high template concentration of 1 µM based on previous optimization to maximize the RNA yield when all rNTPs are in excess (10). However, when UTP was at rate-limiting concentration, lowering of the template concentration to below 150 nM increased the RNA yield more than 3-fold (Figure 1C). The template concentration of 20 nM was considered optimal. Optimization of the BI concentration led to a further increase in sensitivity (Figure 1D). Together these changes enabled the meaningful quantification of UTP (Figures 1E and F).

**Figure 1.**
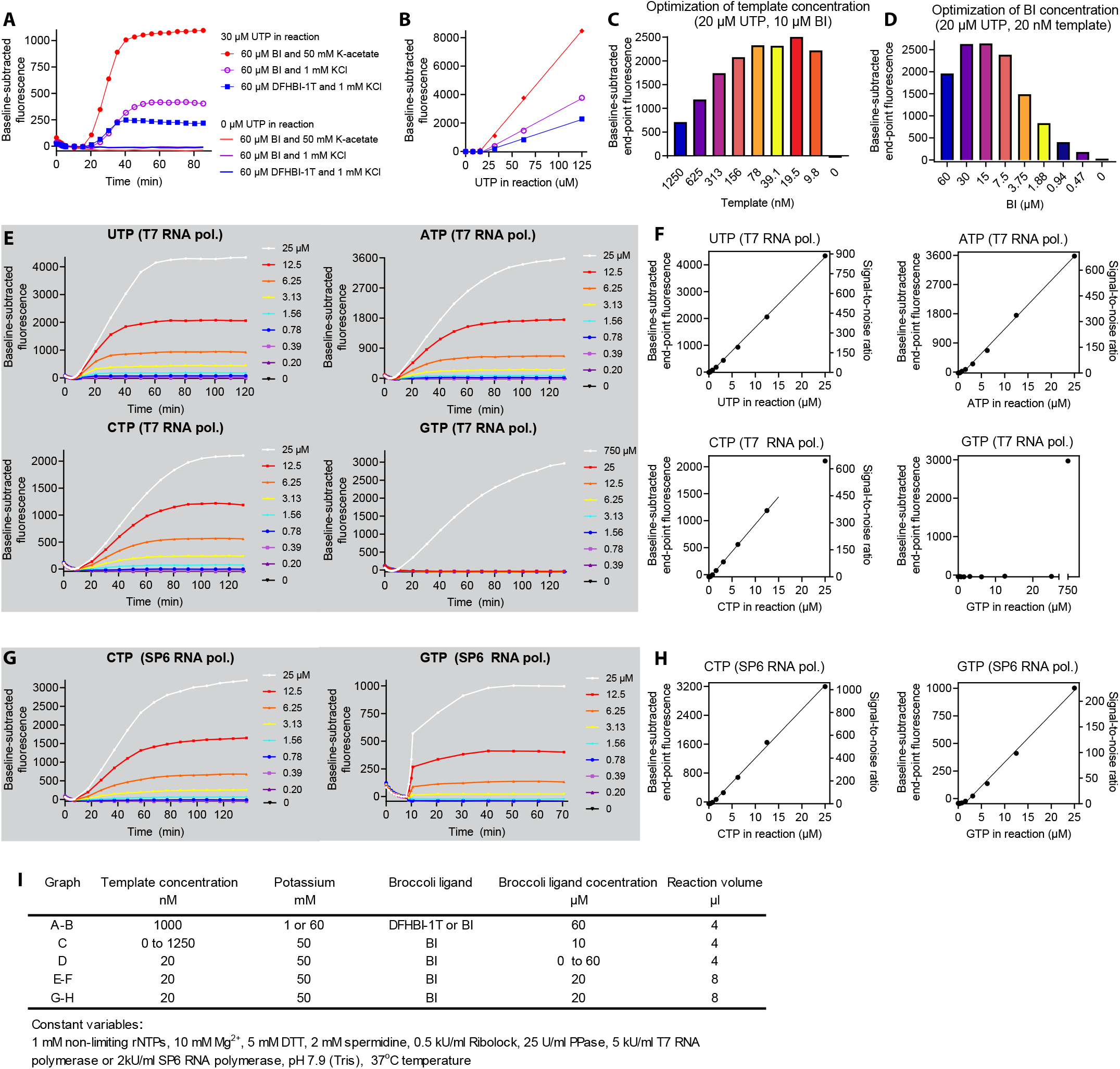
Broccoli aptamer, its fluorogenic ligand BI, and potassium acetate -supplemented buffer enable determination of rNTPs by quantitative *in vitro* transcription. (**A-B**) Comparison of Broccoli ligands (DFHBI-1T and BI) and buffer potassium concentration (enhancer of the aptamer folding) on real-time monitoring of *in vitro* transcription under limiting UTP concentrations with T7 RNA polymerase. (**A**) Representative fluorescence traces recorded during the *in vitro* transcription. (**B**) Standard curves generated from the baseline-subtracted end-point fluorescence values. The symbols and colors represent the same assay conditions as in Figure 1A. (**C**) Optimization of the template concentration. (**D**) Optimization of BI concentration. (**E**) Real-time detection of Broccoli-BI fluorescence during the assay reactions with T7 RNA polymerase, and (**F**) Standard curves generated from the end-point fluorescence values. A linear regression line is shown for the linear part of the standard curve. Real-time detection of Broccoli-BI fluorescence during the CTP and GTP assay reactions with SP6 RNA polymerase, and the corresponding standard curves. (**I**) Assay conditions in graphs A to H.

Next, we tested the assay for the quantification of the other rNTPs. The quantification of ATP and CTP did not require further optimization (Figures 1E and F). However, the quantification of GTP turned out to be problematic with T7 RNA polymerase as it prefers to initiate transcription with three consecutive guanosines (16) (Table 1 and Figures 1E and F). When GTP is severely limiting, T7 RNA polymerase presumably undergoes excessive cycles of abortive initiation, leading to the consumption of GTP without production of full-length transcripts. To overcome this issue, we tested an adenosine-initiating template but without success (Supplementary Figure S1A). SP6 RNA polymerase also prefers to initiate transcription with a guanosine but has less preference for the immediate downstream nucleotide sequence than the T7 RNA polymerase (17). We designed for the SP6 RNA polymerase a template in which a 10-nt pre-aptamer leader sequence contains a single starting guanosine and the next guanosine is at position +11 (Template 2, Table 1). This template and RNA polymerase combination enabled useful dynamic range for the quantification of GTP (Figure 1G and H). SP6 RNA polymerase and template 2 also slightly increased the sensitivity for the quantification of CTP (Figure 1H).

### Further optimization of the assay

After a successful proof-of-concept, we set out to optimize other assay parameters. The optimal template concentration for T7 RNA polymerase (Figure 1C) proved to be optimal for SP6 RNA polymerase as well (Supplementary Figure S1B). Over 15-fold differences in RNA polymerase concentrations had surprisingly little effect on the final aptamer yield when the reactions were allowed to progress to completion (Figures 2A-D). Based on maximal end-point fluorescence within 2h, we selected 1.9 kU/ml as optimal concentration for the T7 RNA polymerase. For SP6 RNA polymerase, we considered 1 kU /ml and 0.5 kU/ml as optimal concentrations for the quantification of CTP and GTP, respectively.

**Figure 2.**
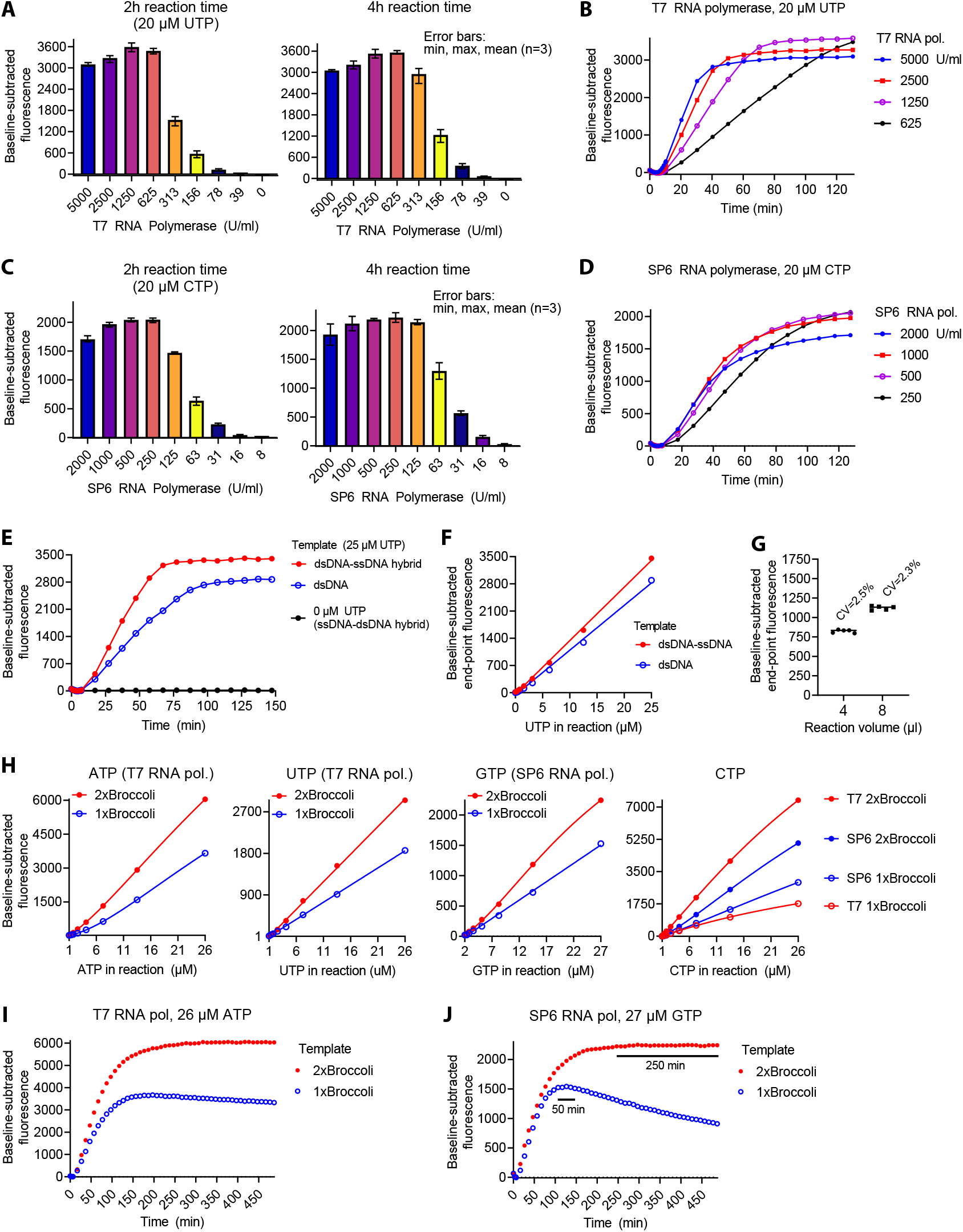
Optimization of *in vitro* transcription for quantification of rNTPs. (**A**-**D**) Optimization of RNA polymerase concentration: T7 RNA polymerase and limiting concentration of UTP (**A**-**B**), SP6 RNA polymerase and limiting concentration of CTP (**C**-**D**). (**E**-**F**) Comparison of double-stranded single-stranded hybrid template and the fully double-stranded equivalent with T7 RNA polymerase. (**G**) Reaction volume and variation of technical replicates (T7 RNA polymerase and 6.25 µM UTP). CV, coefficient of variation. (**H**) Signal amplification with stabilized dimeric Broccoli. UTP quantification reactions were performed in a qPCR instrument, and the rest of the reactions in a plate reader. (**I-J**) Representative fluorescence traces of ATP (**I**) and GTP (**J**) quantification reactions with templates encoding monomeric or dimeric Broccoli. Stable plateau phases are marked with black bar for GTP quantification reactions. The reaction volumes were 10 µl in **H-J** and 4 µl elsewhere if not otherwise mentioned in the figure.

*In vitro* transcription is usually performed at 37-42°C, while the folding of the Broccoli-aptamer is optimal at lower temperatures (15). Decrease of the reaction temperature from 37°C to 30°C slowed down the reaction phase but the end-point fluorescence values were practically identical (Supplementary Figures S1C-H). Cooling down the plateaued reactions to 23°C marginally increased the aptamer fluorescence (Supplementary Figures S1E-H), indicating close to 100% efficient folding of the Broccoli at 37°C under the given conditions.

T7 RNA polymerase requires a double-stranded promoter but efficiently transcribes single-stranded downstream sequence (16). For the experiments above, we used a double-stranded single-stranded hybrid template for the T7 RNA polymerase. A fully double-stranded equivalent proved less optimal (Figures 2E, F). SP6 RNA polymerase has an absolute requirement for fully double-stranded DNA (18). Hence, we did not test hybrid templates with this RNA polymerase.

A T7 RNA polymerase harbouring a P266L point mutation shows enhanced promoter clearance and attenuated abortive initiation (19). We tested these potentially beneficial features with UTP as the rNTP to be quantified. The *in vitro* transcription progressed faster with this mutant than with the wild-type enzyme (Supplementary Figure S1I). However, sensitivity of the assay slightly decreased (Supplementary Figure S1J).

A 384-well PCR plate allows miniaturization of e.g. qPCR reactions to a few microliters. We tested reaction volumes of 4 and 8 µl side by side. The coefficient of variation (CV) between technical replicates was less than 2.6% with both reaction volumes (Figure 2G). The higher reaction volume led to somewhat higher baseline-corrected end-point fluorescence values.

The standard curves for all rNTPs showed wide sensitive linear ranges but clear threshold concentrations below which linear curve fitting did not apply and the resolution of the assay became poor. To eliminate this useless concentration range, we decided to include low basal concentration (1 to 2 µM) of the limiting rNTP in the assay reactions. Due to this modification any increase over the basal concentration led to a steep linear increase in the measured signal (Supplementary Figure S2). Moreover, the linearization of the standard curve enables easy application of the Standard Addition Method to correct matrix effects if required.

Because rNTPs are abundant metabolites and because of the clear threshold concentration (0.2-0.5 µM) of our assay, we did not seek to benchmark the assay based on the traditional lowest limit of detection and quantification. Instead, we concentrated on a more relevant parameter, the quantitative resolution (20), in other words, the lowest analyte concentration difference the assay can discriminate. Using technical triplicates and 8 µl reaction volume, we could reliably quantify rNTP differences less than 4% (Supplementary Figures S1K and L), meaning that the resolution was close to pipetting precision.

As a potential further signal enhancement, we tested templates encoding dimeric Broccoli (Table 1). Instead of concatemeric Broccoli constructs with relative lengthy scaffolds developed for live cell imaging (14), we chose a dimeric Broccoli with a 4 base pair stabilizing stem that prevents misfolding of the aptamer (21). In comparison to the monomeric aptamer, the dimeric counterpart led to approximately 1.6-fold increase in the steepness of ATP, UTP, and GTP standard curves (Figure 2H). In the case of CTP quantification, the benefits of dimeric Broccoli were RNA polymerase and template dependent. While SP6 RNA polymerase provided higher sensitivity than T7 RNA polymerase with the monomeric Broccoli, the reverse was true with dimeric Broccoli. With T7 RNA polymerase the dimeric Broccoli led to over 4-fold increase in the CTP standard curve steepness, making the assay the most sensitive for quantification of CTP (Figure 2H), the least abundant rNTP in many cell types (2). Two drawbacks of the dimeric Broccoli were increased reaction time and necessity to perform non-linear curve fitting for CTP and GTP quantifications (Figures 2H-J). On the other hand, the dimeric Broccoli led to stabilization of the plateau phase of the GTP quantification reactions (Figure 2J). It is possible that this stabilized aptamer construct is resistant to single-strand RNA/DNA exonuclease activity of some RNA polymerases upon GTP exhaustion (22).

### Modification of the assay for rNMPs and rNDPs

Because most biological processes are sensitive to the energy charge of nucleotides (mainly rNTP-to-rNDP ratio), knowledge on the concentrations of rNMPs and rNDPs can be as important as that of rNTPs. We asked whether we could modify our assay for these nucleotide species by phosphorylating the mono- and diphophates to triphosphates. First, we added a nucleoside diphosphate kinase in the assay reactions and measured the rNTP signal from samples with equal total nucleotide concentration but varying proportions of rNTPs and rNDPs. In this approach, the nucleoside diphosphate kinase utilizes the non-limiting rNTPs as phosphate donors to phosphorylate the target rNDP to the corresponding rNTP, allowing the measurement of the sum of rNDP+rNTP of interest. All rNDP+rNTP mixtures gave essentially the same expected rNTP signal (Figure 3A), implying essentially complete rNDP-to-rNTP phosphorylation. Monitoring of the transcription kinetics did not reveal differences between rNDP or rNTP samples or their mixtures, indicating almost instant rNDP phosphorylation (Figure 3B).

**Figure 3.**
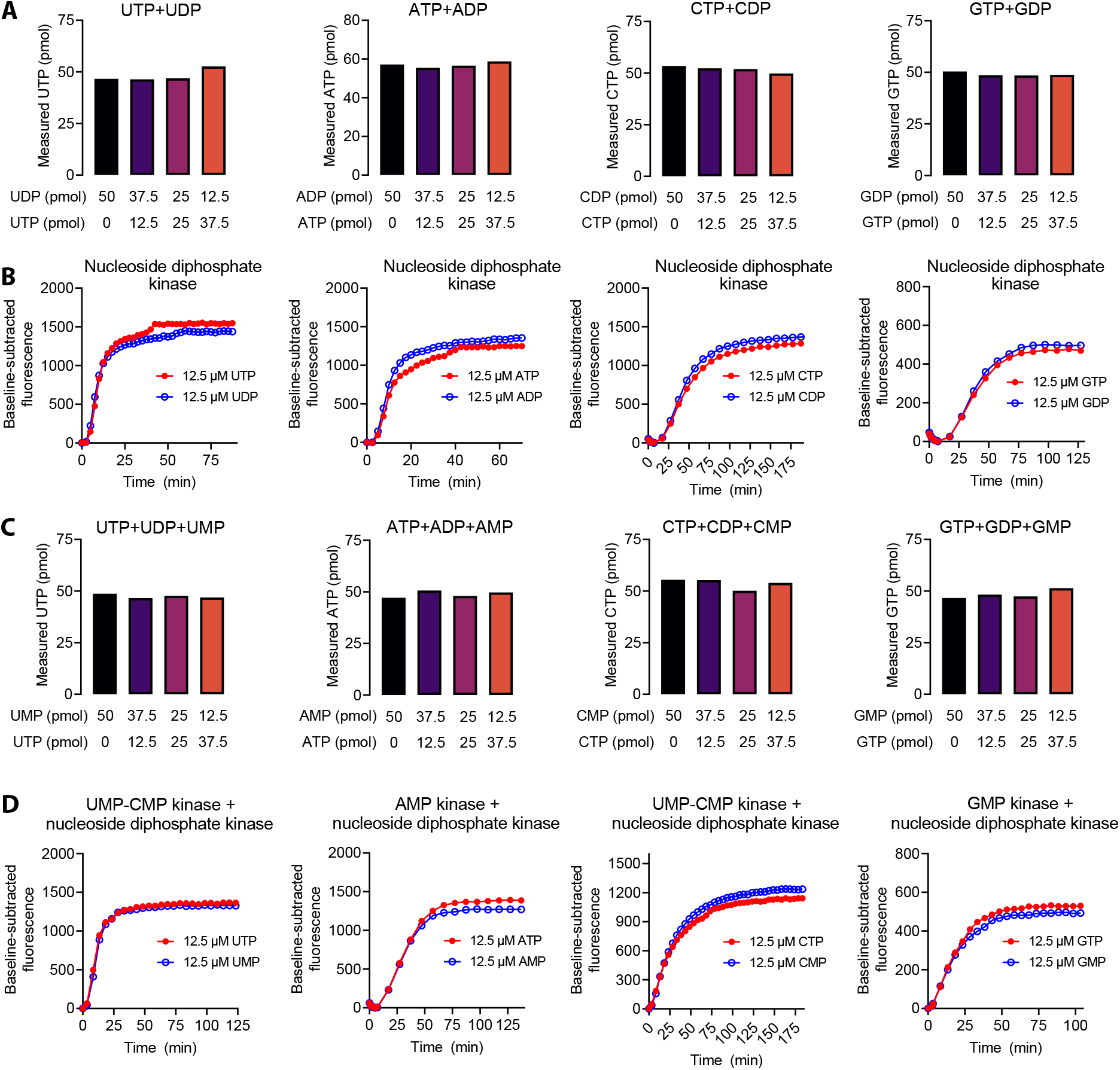
Quantification of rNDPs and rNTPs. (**A**) Quantification of the indicated rNDP and rNTP combinations by inclusion of nucleoside diphosphate kinase in the assay reactions. (**B**) Representative fluorescence traces of assay reactions containing nucleoside diphosphate kinase and either rNTP or rNDP as a limiting nucleotide. (**C**) Quantification of the indicated rNMP and rNTP combinations by inclusion of nucleoside diphosphate kinase and rNMP kinases in the assay reactions. (**D**) Representative fluorescence traces of assay reactions containing nucleoside diphosphate kinase and rNMP kinase and either rNTP or rNMP as a limiting nucleotide.

**Figure 4.**
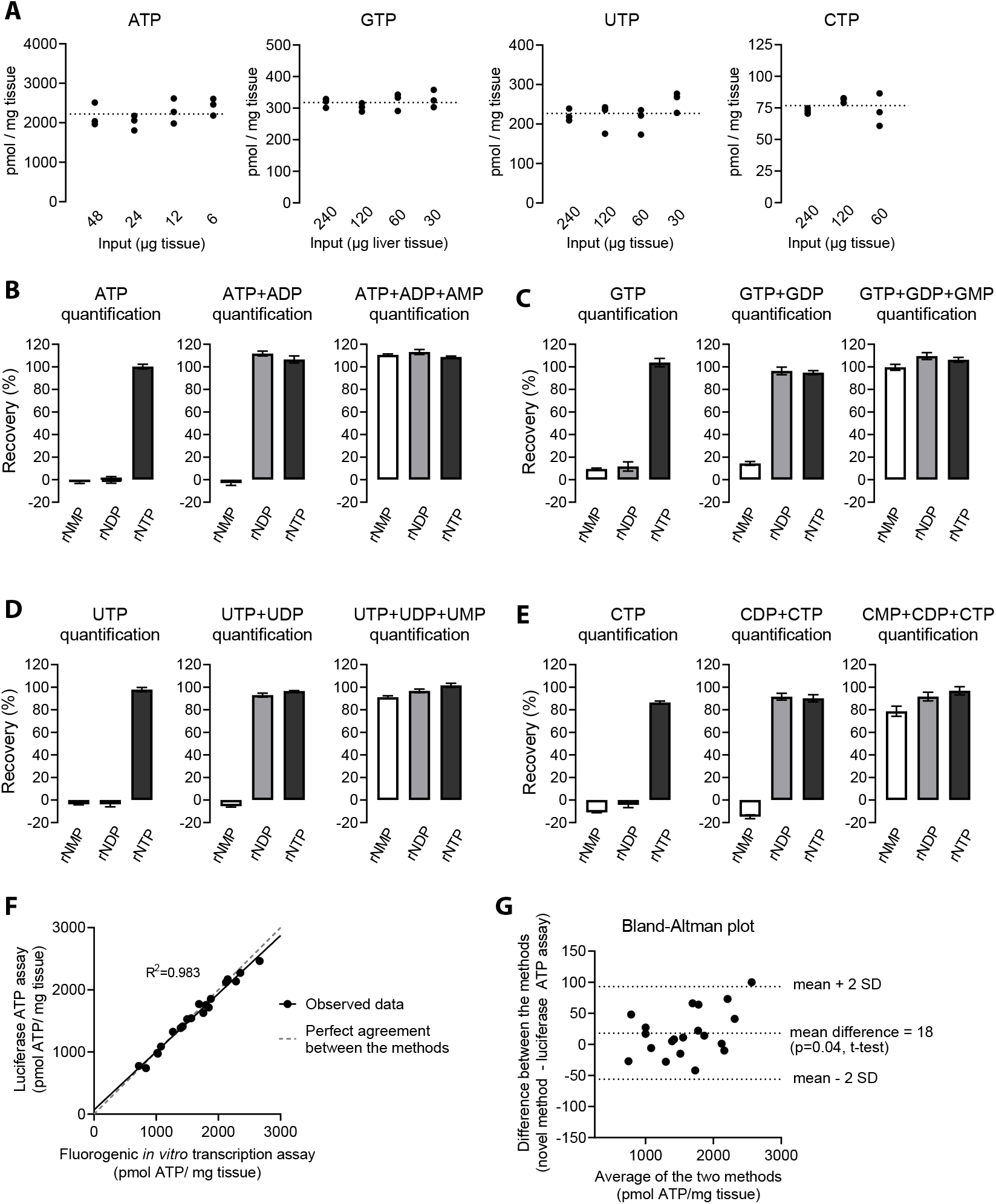
Quantification of rNTPs from mouse liver. (**A**) Estimation of sample-related interference by dilution series. The input refers to polar metabolite extract worth of initial tissue mass. The reaction volumes were 4 µl (1 µl sample + 3 µl assay reagent). The data points are technical replicates from 4 pooled liver extracts. (**B**-**E**) Recovery of exogenous rNMPs, rNDPs, and rNTPs spiked into liver extracts. The purified nucleotides were added as mixtures comprising 2000 pmol adenosine nucleotides (AMP, ADP, or AMP), 200 pmol uridine nucleotides (UMP, UDP, or UTP), 200 pmol guanosine nucleotides (GMP, GDP, or GTP), and 50 pmol cytidine nucleotides (CMP, CDP, or CTP) per mg tissue. Sample inputs (relative to pre-extraction tissue weight) were following: 11.25 µg for ATP, 112.5 µg for GTP and UTP, and 450 µg for CTP. The reaction volumes were 10 µl (5 µl sample + 5 µl assay reagent). The bar graphs represent mean and SEM from 4 independent liver extracts. (**F**) Correlation between hepatic ATP levels as determined by the novel method and luficerin-luciferase ATP assay. The variable ATP levels were achieved by warm ischemia (Supplementary Figure 4). (**G)** Bland-Altman plot visualization of the differences between the novel method and the luficerin-luciferase ATP assay.

Next, we added the nucleoside diphosphate kinase and AMP kinase, or GMP kinase, or UMP-CMP kinase in the assay reactions to determine the rNMPs. We repeated the previous experiment but now with varying mixtures of rNMPs and rNTPs. Again, the phosphorylation to rNTPs was almost instant and all mixtures gave essentially the same expected rNTP signal (Figures 3C and D).

### Application of the assay to biological samples

We extracted polar metabolites from mouse liver tissue and made a dilution series of the extracts to assess the applicability of the assay for complex biological samples and proneness to sample-related interference. Each of the four canonical rNTPs were easily quantifiable with our assay (Figure 4A). All dilutions gave essentially the same tissue rNTP concentrations, excluding major sample-related inhibition. The measured liver rNTP concentrations were very close to the values reported for rat liver in the systematic review of Traut el al. (2). As an alternative approach to assess sample-related interference, we spiked known amounts of rNTPs to liver extracts and assessed the *in vitro* transcription efficiency in comparison to purified rNTPs. The recovery of ATP, GTP, and UTP spiked into the extracts was essentially 100% (Figures 4B-D). The recovery of CTP was slightly less (range: 84-96%) (Figure 4E, Supplementary Figure S3). We also assessed liver extracts spiked with rNMPs and rNDPs. We found that extracts that had not undergone heat denaturation step showed rNMP kinase activity that could phosphorylate at least CMP (Supplementary Figure S3). Slight modifications to the extraction protocol to increase the stringency of extract deproteination, including heat denaturation step, completely abolished this interference (Figure 4E, Materials and Methods). After these modifications, no unintended nucleotide interconversion took place (Figure 4B-E). With addition of exogenous nucleotide kinases to quantify rNMPs and rNDPs, the recoveries of these nucleotides spiked to liver extracts were similar to that of rNTPs (Figures 4B-E).

To further test the utility of our assay, we performed a classic warm ischemia experiment to disrupt nucleotide pools *ex vivo*, exposing subsamples from mouse livers to periods at room temperature (Figure 5, Supplementary Figure S4). The ATP levels steeply declined for the first 4 minutes of the ischemia and then briefly stabilized before the decline continued (Figure 5A). After 10 minutes, the ATP levels were 39% of the initial values. ADP levels were relatively stable (Figure 5B). In contrast, AMP immediately started to accumulate, and its levels mirrored the decline of ATP (Figure 5C). The adenylate energy charge dropped from 0.74 to 0.47 during the follow up of 10 minutes (Figure 5D). At the same time, total adenosine nucleotide content gradually and linearly decreased (Figure 5E).

**Figure 5.**
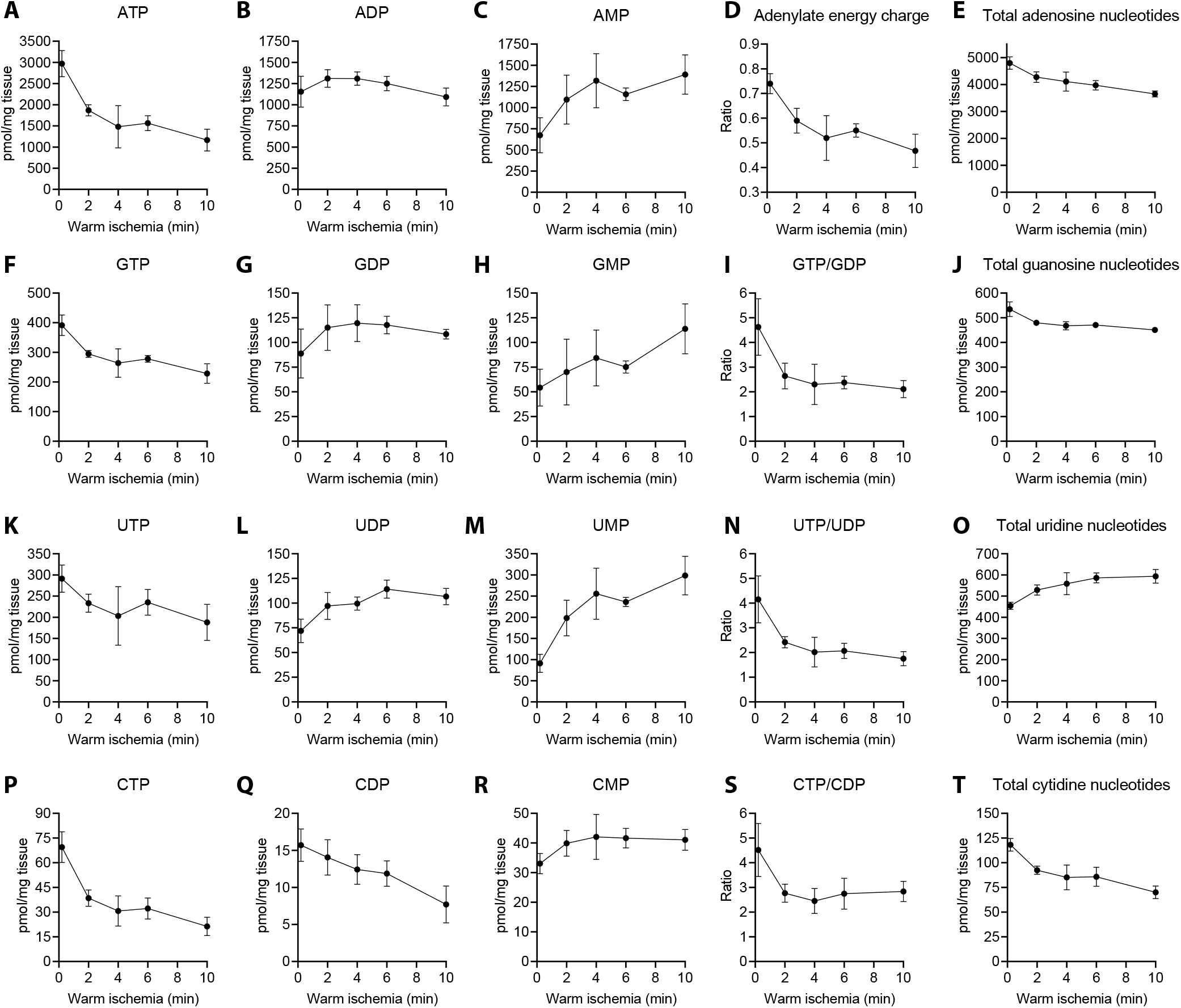
Effect of warm ischemia on hepatic ribonucleotide pools. (**A-C**) Quantification of adenylate nucleotides. (**D**) Adenylate energy charge defined as (ATP+0.5ADP)/(ATP+ADP+AMP). (**E**) Total adenylate nucleotides (ATP+ADP+AMP). (**F**-**J**) Quantification of guanidine nucleotides, and GTP/GDP ratio. (**K**-**O**) Quantification of uridine nucleotides and UTP/UDP ratio. (**P**-**T**) Quantification of cytidine nucleotides and CTP/CDP ratio. The data points represent mean and SD from four non-fasted male mice of 2 months of age. Where error bars are not visible, the SD was smaller than the size of the symbol. Supplementary Figure 4 shows data from a similar experiment in which the extracts were prepared as described in Supplementary methods (no boiling step).

As a reference method, we employed luciferin-luciferase enzymatic assay to remeasure ATP from the ischemic liver samples. The correlation coefficient between the methods was 0.983, and the mean difference across the range of 750 to 2600 pmol ATP/mg liver tissue was 18 pmol (Figures 4F and G).

Our assay was also able to quantify all other canonical ribonucleotides and their disturbances during the ischemia. GTP and UTP showed similar patterns of ischemic decline to ATP but with less steep changes (Figures 5F and K). CTP showed the steepest decline among the four rNTPs (Figure 5P). In contrast to the rather stable ADP levels (Figure 5C), GDP and UDP clearly accumulated during the ischemia (Figures 5G and L), leading to a clear decrease in rNTP-to-rNDP ratio of the respective nucleotides (Figure 5I,N). Instead, CDP levels relatively linearly decreased while CTP-to-CDP ratio still collapsed due even greater loss of CTP (Figures 5P, Q and S). GMP levels were somewhat variable but showed a clear pattern of accumulation (Figure 5H). UMP accumulated relatively even more than AMP (Figures 5C and M), while CMP showed a small acute increase during the first few minutes and then stayed stable (Figure 5R). The size of the total cytidine nucleotide pool decreased linearly and was roughly 50% of the baseline level after 10 minutes (Figure 5T). The total guanosine pool also linearly decreased but more modestly, while total uridine nucleotide pool actually increased during the ischemia (Figure 5J,O), potentially due to degradation of nucleotide sugars. Taking into account potential species-specific differences (rat vs mouse), differences between fasted and non-fasted animals, and differing methodology related to sample processing, our results were highly in line with rodent liver reference data (2, 23).

Finally, we assessed the impact of disrupted oxidative phosphorylation on nucleotide levels in two cell lines originating from mouse liver. In non-transformed AML12 cell line, which derives from hepatocytes of mice overexpressing human transforming growth factor α (24), four inhibitors targeting different sites/processes of the oxidative phosphorylation collapsed the levels of all rNTPs and adenylate energy charge during a 6-hour exposure (Figure 6A), indicating heavy reliance of these cells on mitochondria for maintenance of nucleotide pools, similar to observed in primary hepatocytes (25). In contrast, the adenylate energy charge and purine nucleotide concentrations were totally indifferent to blockade of oxidative phosphorylation in Hepa 1-6 hepatoma cells (Figure 6B). Instead, myxothiazol, which disrupts the oxidative phosphorylation at the level of cytochrome *bc*_1_ complex (respiratory complex III), caused a selective pyrimidine nucleotide depletion in these cells. The pyrimidine nucleotide biosynthesis connects to mitochondrial respiration at the level of dihydroorotate dehydrogenase (DHODH), which utilizes ubiquinone as the electron acceptor (26). The function of cytochrome *bc*_1_ complex is to oxidize ubiquinol to ubiquinone, and therefore many cancer cells, and apparently Hepa 1-6 cells as well, depend on mitochondrial respiration for pyrimidine nucleotide biosynthesis but not necessary for purine nucleotide biosynthesis and ATP generation (27).

**Figure 6.**
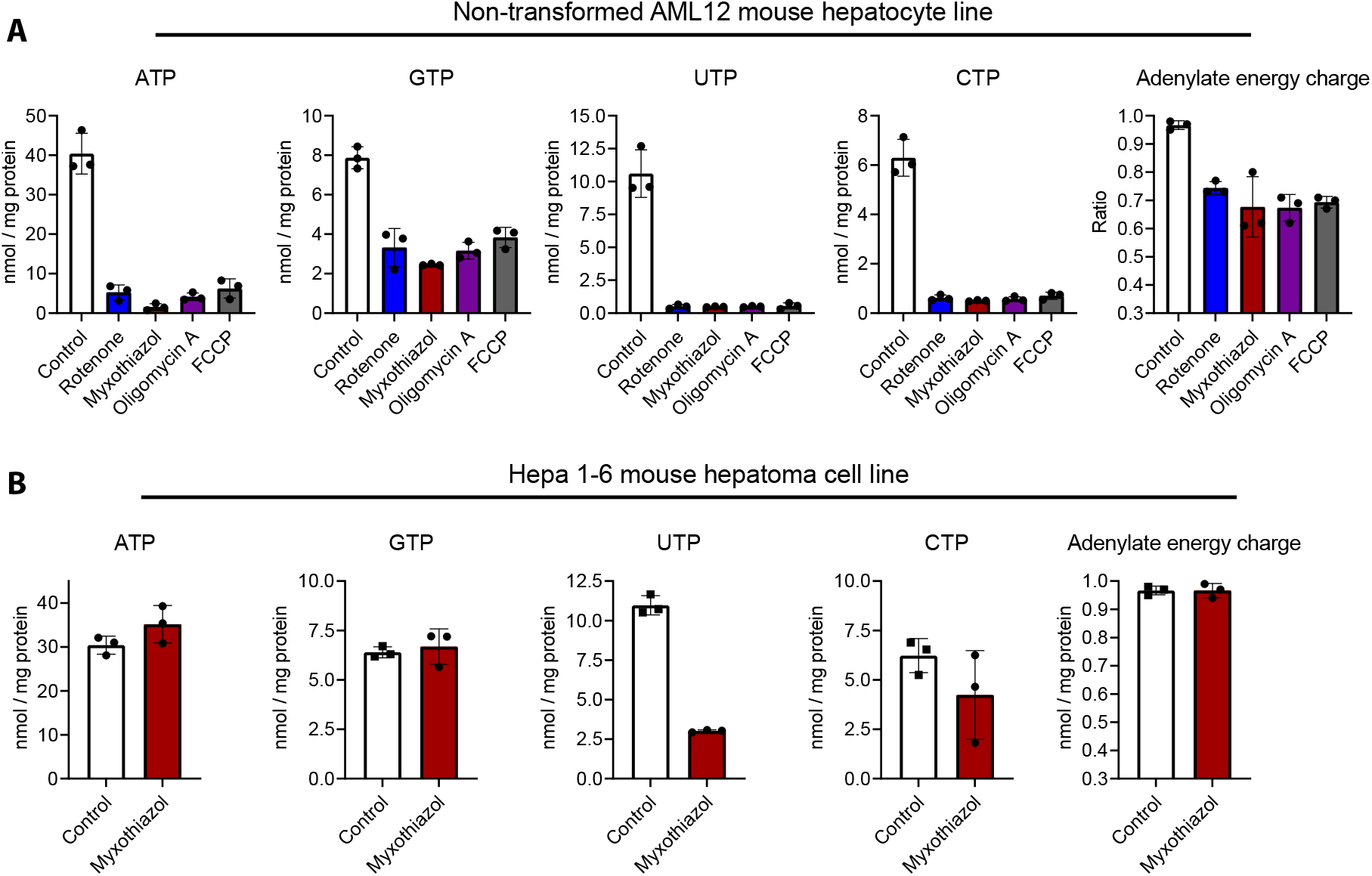
rNTP levels and adenylate energy charge after inhibition of oxidative phosphorylation in cultured mammalian cells. (**A**) Data from AML12 mouse hepatocyte line. (**B**) Data from Hepa 1-6 mouse hepatoma cell line. The inhibitor concentrations were: 100 nM rotenone, 100 nM myxothiazol, 1 µM oligomycin A, and 5 µM FCCP. These inhibitors target mitochondrial NADH dehydrogenase (complex I), cytochrome *bc*_1_ complex (complex III), ATP synthase, and mitochondrial membrane potential, respectively. The data points represent individual cell culture dishes and the error bars SD.

### Precision of the assay

Table 2 and 3 show typical CVs for rNTP standards and liver extracts, respectively. Generally, the CV values were well below 2.5%.

**Table 2.** Intra-assay coefficient of variation (CV) of rNTP standard samples

| rNTP | Input (pmol) | Mean (%) | SD (%) |
| --- | --- | --- | --- |
| ATP | 125 | 0.9 | 0.2 |
| ATP | 62.5 | 1.4 | 1.0 |
| ATP | 31.3 | 1.5 | 0.4 |
| ATP | 15.6 | 2.8 | 1.5 |
| ATP | 7.81 | 4.5 | 0.6 |
| GTP | 125 | 1.5 | 0.6 |
| GTP | 62.5 | 2.3 | 1.2 |
| GTP | 31.3 | 1.8 | 1.6 |
| GTP | 15.6 | 1.6 | 0.5 |
| GTP | 7.81 | 5.6 | 2.2 |
| UTP | 125 | 1.4 | 1.1 |
| UTP | 62.5 | 1.8 | 0.6 |
| UTP | 31.3 | 3.5 | 4.0 |
| UTP | 15.6 | 2.4 | 1.5 |
| UTP | 7.81 | 2.4 | 2.5 |
| CTP | 125 | 2.5 | 1.7 |
| CTP | 62.5 | 1.9 | 1.6 |
| CTP | 31.3 | 1.2 | 0.5 |
| CTP | 15.6 | 1.4 | 1.1 |
| CTP | 7.81 | 1.5 | 0.5 |
| CTP | 3.91 | 3.6 | 2.4 |

The mean and SD represent CVs from three independent triplicate measurements. The reaction volumes were 10 μl (5 μl sample and 5 μl assay reagent).

**Table 3.** Intra-assay coefficient of variation (CV) of liver extracts

| rNTP | Mean (%) | SD (%) |
| --- | --- | --- |
| ATP | 1.7 | 0.9 |
| ATP+ADP | 1.3 | 0.7 |
| ATP+ADP+AMP | 1.3 | 0.5 |
| GTP | 1.9 | 0.7 |
| GTP+GDP | 1.7 | 1.1 |
| GTP+GDP+GMP | 1.2 | 0.8 |
| UTP | 1.5 | 0.9 |
| UTP+UDP | 1.2 | 0.6 |
| UTP+UDP+UMP | 1.4 | 0.8 |
| CTP | 1.7 | 0.9 |
| CTP+CDP | 1.5 | 0.9 |
| CTP+CDP+CMP | 1.5 | 1.0 |

## DISCUSSION

The bacteriophage T7 and SP6 RNA polymerases are extensively characterized enzymes and vital molecular biology tools. A few studies have also investigated the enzymological properties of these two RNA polymerases under limiting nucleotide concentrations (9, 28, 29), but the current study demonstrates for the first time their successful employment for the quantification of rNTPs. The assay we describe is simple and straightforward requiring only mixing a detection reagent and a sample in a microplate well. A typical qPCR instrument or plate reader can be used to read the plate and all the reagents are commercially available. The assay shows very low intrinsic variation, meaning that technical replicates are not an absolute requirement for precise rNTP quantification with a good pipetting technique. The modifications enabling the determination of rNMPs and rNDPs, make the assay a highly versatile tool to essentially any laboratory in a need for a convenient method to quantify ribonucleotides.

While methods involving anion-exchange or hydrophilic interaction liquid chromatography, or capillary electrophoresis exist (4), many research groups do not have access to the expensive specialized equipment required by these methods. Moreover, many established chromatography procedures are poorly compatible with modern equipment coupled to mass spectrometer, leading to additional challenges in the quantification of nucleotides. In fact, a motivation for this study was several failed attempts to measure UTP, UDP, and UMP by ultra-performance liquid chromatography-mass spectrometry.

We optimized several parameters relevant to the novel assay but found the optimization of the *in vitro* transcription conditions *per se* unnecessary because of the extensive literature on the topic (10, 30). We, however, do acknowledge the following points as potentially worthy of further assay development. We chose the Broccoli aptamer as the fluorescent reporter. The main rationale for this decision was its short length (49 nt) and it being well characterized (12, 15), including for real-time monitoring of *in vitro* transcription (10). Moreover, the recently identified Broccoli ligand BI makes this aptamer exceptionally stable at optimal *in vitro* transcription temperatures (14). In comparison to the original and more widely used ligand the DFHBI-1T, BI also induces brighter fluorescence. Numerous other fluorescent aptamer-ligand pairs do, however, exist and may possess potentially beneficial features for the rNTP quantification (recently reviewed in (11)). Because abortive initiation is the main cause of truncated unproductive RNA, it may be possible to increase the sensitivity of the assay by designing templates that encode several concatenated aptamers. A similar approach has facilitated imaging of single transcripts in live cells (14). Our experiments with dimeric Broccoli, however, suggest that further aptamer concatamerization likely provides only incremental benefits. We settled for two templates that allow a robust quantification of the four canonical rNTPs. The most optimal template, however, depends on the rNTP species to be quantified. There is not much room for the manipulation of the promoter or the aptamer sequences, but the pre-aptamer leader sequences are adjustable for minimization of abortive initiation and enhanced promoter clearance to increase the sensitivity of the assay. By employing a suitable set of mono- and dinucleoside kinases, we extended the assay to cover all canonical ribonucleotide species. In theory, a similar set of kinases targeting the unphosphorylated ribonucleosides would enable determination of adenosine, uridine, cytidine, and thymidine as well. A parallel quantification of dNTPs is also possible with our previously published method that also has similarly convenient real-time fluorometric detection in qPCR plate format (5).

We demonstrated the utility of our method by measuring the ribonucleotides from mouse liver tissue. We excluded major sample-related interference with this tissue and the given extract dilutions. If, nonetheless, interference is encountered and absolute quantification is required, our method is amenable to the classic Standard Addition Method as a correction. More likely source of error is, however, any delay in the quenching of enzymatic activity during sample collection as our ischemia time series and several other studies demonstrate (23, 31, 32). As the final readout in our method is the increase of fluorescence over the baseline, a modest amount of sample-derived intrinsic fluorescence is not generally a significant source of error. Given the nature of the assay, it can be considered inherently specific as unspecific RNA synthesis is expected to compromise the aptamer structure. Some unnatural RNA-incorporating nucleotide analogues can potentially, however, give signal with our method. More relevantly, however, it is important to follow stringent-enough extraction protocol to avoid sample-derived enzymatic activity and unintended nucleotide interconversion. The rNTP concentrations in the liver as measured by our method were well within published high-quality reference values (2). Moreover, we were able to almost exactly replicate the classic experiment of the effect of ischemia on nucleotide pools (23), validating further the applicability of the assay for determination of not only rNTPs but rNMPs and rNDPs as well.

Targeting nucleotide metabolism is increasingly studied as a therapy for cancer, viral infections, and inflammatory diseases (33). Moreover, compromised biosynthesis and maintenance of phosphorylation statuses of nucleotides are proposed primary disease mechanisms in many metabolic diseases such mitochondrial disorders, yet currently reports of alterations in ribonucleotide levels in these diseases and their model organisms are surprisingly scarce. Given the ubiquitous role of nucleotides in cellular processes, the assay reported here is expected to facilitate research in the various fields of life sciences.

## Supporting information

Step-by-step protocol - Quantification of ribonucleotides

Step-by-step protocol - sample extraction

## ACKNOWLEDGEMENTS

We thank Vilma Wanne and Diviya Upadhyay for technical assistance, and Nina Sipari and the Viikki metabolomics unit (Helsinki Institute of Life Science) for the efforts to quantify uridine nucleotides by UPLC-MS, and Prof. Vineta Fellman for critically reading the manuscript. This study was supported by funding from Samfundet Folkhälsan, Jane and Aatos Erkko Foundation, and Medicinska understödsföreningen Liv och Hälsa. Part of the work was carried out with the support of HiLIFE Laboratory Animal Center Core Facility, University of Helsinki, Finland.

