## Supplementary material for "Quantification of all 12 canonical ribonucleotides by real-time fluorogenic *in vitro* transcription": Step-by-step protocol - Quantification of ribonucleotides

(Purhonen and Kallijärvi, 2023)

**Common reagents and materials for all quantifications**

Stock nucleotide solutions and standards should be stored at -80°C. All other reagents should be stored at -20°C. Use molecular biology-grade water and DMSO, and filter tips for dilutions and preparation of the reagents.

- 10x CutSmart buffer (New England BioLabs)
- Molecular biology-grade 100 mM rNTP stocks (e.g. N0466, New England BioLabs). For long-term storage, keep these solutions at 80°C (short-term storage at -20°C is possible).
- 10x non-limiting rNTP mixes [Note 1].

  **C**+**G**+**U** mix for ATP quantification
  10 mM concentration of each non-limiting rNTP (CTP+GTP+UTP) + 10 µM ATP

  **A**+**C**+**G** mix for UTP quantification
  10 mM concentration of each non-limiting rNTP (ATP+CTP+GTP) + 10 µM UTP

  **A+G+U** mix for CTP quantification
  10 mM concentration of each non-limiting rNTP (ATP+GTP+UTP) + 10 µM CTP

  **A+C+U** mix for GTP quantification
  10 mM concentration of each non-limiting rNTP (ATP+CTP+UTP) + 20 µM GTP.

- 6.37 M Spermidine (Sigma-Aldrich, S2626). Equilibrate to room temperature before opening the vial. Dilute an 80 mM working solution in water.
- DL-Dithiothreitol (DTT) (Sigma-Aldrich, D0632).
  100 mM DTT = 15.4 mg/ml water.
- BI dihydrochloride (Tocris Biotechne, Cat. No. 7466, MW= 441.26 g/mol)
  50 mM stock = 22.06 mg/ml DMSO
  Working solution = 1 mM in DMSO.
- 40 kU/ml RiboLock RNAse inhibitor (Thermo, EO0382).
- 100 U/ml Inorganic pyrophosphatase (PPase) (Thermo #EF0221)
  Dilute the 100 U/ml enzyme 1:10 with the provided storage buffer to prepare 10 U/ml working solution.
- qPCR plate (e.g. Bio-Rad #HSP3805 or #HSP3866)
- qPCR plate sealing film (e.g. Bio-Rad #MSB1001) .

**Reagents for ATP, UTP, and CTP quantification**

Dissolve lyophilized oligonucleotides in water.

- 100 µM T7 RNA promoter strand
  5’-TAATACGACTCACTATAG-3’

- 10 µM T7 Broccoli aptamer antisense strand (encoding stabilized dimeric Broccoli)
  5’-
  GAGGGAGCCCACACTCTACTCGACATCTGAGCCCACACTCTACTCGACAGATACGAATATCTGGACCCGACCGTCTCAGATGGACC
  CGACCGTCTCCCTCCCTATAGTGAGTCGTATTA-3’

- Preparation of 5 µM template stock and 0.8 µM working solution for T7 RNA polymerase:
  a) Preheat heat block to 95°C.
  b) Mix 1 volume of 100 µM of the promoter strand,
     10 volumes of 10 µM of antisense strand
     and 9 volumes of water.
  c) Place the mix into the preheated heat block. Switch off the heat block.
  d) Allow the mixture to cool down slowly to room temperature. Thereafter, keep on ice.
  e) Dilute a 0.8 µM working solution.
  f) Store the stock and the working solution at -20°C.

- 50 kU/ml T7 RNA polymerase (New England BioLabs, #M0251)

**Reagents for CTP and GTP quantification**

- 10 µM SP6_Broccoli-SP6_sense
  5’-
  ATTTAGGTGACACTATAGAATACAAAAGAGGGAGACGGTCGGGTCCATCTGAGACGGTCGGGTCCAGATATTCGTATCTGTCGAGT
  AGAGTGTGGGCTCAGATGTCGAGTAGAGTGTGGGCTCCCTC-3’
- 10 µM SP6_Broccoli-SP6_antisense
  5’-
  GAGGGAGCCCACACTCTACTCGACATCTGAGCCCACACTCTACTCGACAGATACGAATATCTGGACCCGACCGTCTCAGATGGACC
  CGACCGTCTCCCTCTTTTGTATTCTATAGTGTCACCTAAAT-3’

- Preparation of 5 µM template stock and 0.8 µM working solution for SP6 RNA polymerase:
  a) Preheat heat block to 95°C.
  b) Mix 1 volume of 10 µM 10 µM SP6_Broccoli-SP6_sense
     and 1 volume of 10 µM SP6_Broccoli-SP6_antisense
  c) Place the mix into the preheated heat block. Switch off the heat block.
  d) Allow the mixture to cool down slowly to room temperature. Thereafter, keep on ice.
  e) Prepare a 0.8 µM working solution.
  f) Store the stock and the working solution at -20°C.

- 20 kU/ml SP6 RNA polymerase (New England Biolabs, #M0207)

**Additional reagents for the quantification of nucleoside mono- and diphosphates**

- **Kinases:**
  0.2 mg/ml AMP Kinase (Bionukleo, NMPK-01), EC:2.7.4.10
  0.2 mg/ml GMP kinase (Bionukleo, NMPK-21), EC: 2.7.4.8
  0.2 mg/ml UMP-CMP kinase (Bionukleo, NMPK-22), EC: 2.7.4.14
  0.2 mg/ml NDP kinase (Bionukleo, NDPK-33), EC: 2.7.4.6

#### Master mixes

The reaction volumes can be scaled up or down to improve pipetting precision or to decrease the cost of the assay, respectively. The master mixes should be kept ice cold and distributed to the wells quickly to prevent warming and premature initiation of the reactions. Without RNA polymerase, the master mixes are relatively stable on ice. The RNA polymerase and kinases should be added just prior to use. The reaction times can be adjusted by increasing or decreasing the RNA polymerase concentration. In general, longer reaction times give higher sensitivity than shorter ones.

- Master mix for **<u>ATP</u>**, **<u>UTP</u>**, or **<u>CTP</u>** quantification
  Recipe for 100 reactions (10 µl final reaction volume: 5 µl master mix and 5 µl sample).
  105 µl $H_2O$
  100 µl 10x NEB CutSmart buffer
  100 µl of 10 mM non-limiting rNTP mix.
  25 µl of 0.8 µM DNA template for T7 RNA pol.
  50 µl of 100 mM DTT
  25 µl of 80 mM spermidine
  20 µl of 1 mM BI
  12.5 µl of 40 kU/ml RiboLock
  25 µl of 10 U/ml PPase
  37.5 µl of 50 kU/ml T7 RNA pol.

Final concentrations (after addition of sample): 1x reaction buffer (20 mM Tris-acetate pH 7.9, 10 mM Mg-acetate, 50 mM potassium acetate, 0.1 mg/ml BSA), 2 mM Spermidine, 1 mM non-limiting rNTPs, 20 nM DNA template, 5 mM DTT, 20 µM BI, 0.5 kU/ml RiboLock 0.25 U/ml PPase, 1.875 kU/ml T7 RNAPol.

- Master mix for **<u>GTP</u>** quantification
  Recipe for 100 reactions (10 µl final reaction volume: 5 µl master mix and 5 µl sample).
  117.5 µl $H_2O$
  100 µl 10x NEB CutSmart buffer
  100 µl of 10 mM non-limiting rNTP mix.
  25 µl of 0.8 µM DNA template for SP6 RNA pol.
  50 µl of 100 mM DTT
  25 µl of 80 mM spermidine
  20 µl of 1 mM BI
  12.5 µl of 40 kU/ml RiboLock
  25 µl of 10 U/ml PPase
  25 µl of 20kU/ml SP6 RNA polymerase

Final concentrations (after addition of sample): 1x reaction buffer (20 mM Tris-acetate pH 7.9, 10 mM Mg-acetate, 50 mM potassium acetate, 0.1 mg/ml BSA), 2 mM Spermidine, 1 mM non-limiting rNTPs, 20 nM DNA template, 5 mM DTT, 20 µM BI, 0.5 kU/ml RiboLock, 0.25 U/ml PPase, 0.5 kU/ml SP6 RNAPol

- Master mix for **ADP+ATP, UDP+UTP,** or **CDP+CTP** quantification
  Recipe for 100 reactions (10 µl final reaction volume: 5 µl master mix and 5 µl sample).
  80 µl $H_2O$
  100 µl 10x NEB CutSmart buffer
  100 µl of 10 mM non-limiting rNTP mix.
  25 µl of 0.8 µM DNA template for T7 RNA pol.
  50 µl of 100 mM DTT
  25 µl of 80 mM spermidine
  20 µl of 1 mM BI
  12.5 µl of 40 kU/ml RiboLock
  25 µl of 10 U/ml PPase
  25 µl of 0.2 mg/ml NDP kinase (BioNukleo, NDPK-33)
  37.5 µl of 50 kU/ml T7 RNA pol (NEB).

Final concentrations (after addition of sample): 1x reaction buffer (20 mM Tris-acetate pH 7.9, 10 mM Mg-acetate, 50 mM potassium acetate, 0.1 mg/ml BSA), 2 mM Spermidine, 1 mM non-limiting rNTPs, 20 nM DNA template, 5 mM DTT, 20 µM BI, 0.5 kU/ml RiboLock, 0.25 U/ml PPase, 5 µg/ml NDP kinase, 1.875 kU/ml T7 RNAPol.

- Master mix for **AMP+ADP+ATP, UMP+UDP+UTP,** or **CMP+CDP+CTP** quantification
  Recipe for 100 reactions (10 µl final reaction volume: 5 µl master mix and 5 µl sample).
  55 µl $H_2O$
  100 µl 10x NEB CutSmart buffer
  100 µl of 10 mM non-limiting rNTP mix.
  25 µl of 0.8 µM DNA template for T7 RNA pol.
  50 µl of 100 mM DTT
  25 µl of 80 mM spermidine
  20 µl of 1 mM BI
  12.5 µl of 40 kU/ml RiboLock
  25 µl of 10 U/ml PPase
  25 µl of 0.2 mg/ml NMP kinase*
  25 µl of 0.2 mg/ml NDP kinase (BioNukleo, NDPK-33)
  37.5 µl of 50 kU/ml T7 RNA pol (NEB)

  *AMP Kinase (Bionukleo, NMPK-01) for AMP quantification
  *UMP-CMP kinase (Bionukleo, NMPK-22) for UMP and CMP quantification

Final concentrations (after addition of sample): 1x reaction buffer (20 mM Tris-acetate pH 7.9, 10 mM Mg-acetate, 50 mM potassium acetate, 0.1 mg/ml BSA), 2 mM Spermidine, 1 mM non-limiting rNTPs, 20 nM DNA template, 5 mM DTT, 20 µM BI, 0.5 kU/ml RiboLock, 0.25 U/ml PPase, 5 µg/ml NMP kinase, 5 µg/ml NDP kinase, 1.875 kU/ml T7 RNAPol.

- Master mix for **GDP+GTP** quantification
  Recipe for 100 reactions (10 µl final reaction volume: 5 µl master mix and 5 µl sample):
  92.5 µl $H_2O$
  100 µl 10x NEB CutSmart buffer
  100 µl of 10 mM non-limiting rNTP mix.
  25 µl of 0.8 µM DNA template for SP6 RNA pol.
  50 µl of 100 mM DTT
  25 µl of 80 mM spermidine
  20 µl of 1 mM BI
  12.5 µl of 40 kU/ml RiboLock
  25 µl of 10 U/ml PPase
  25 µl of 0.2 mg/ml NDP kinase (BioNukleo, NDPK-33)
  25 µl of 20kU/ml SP6 RNA polymerase

Final concentrations (after addition of sample): 1x reaction buffer (20 mM Tris-acetate pH 7.9, 10 mM Mg-acetate, 50 mM potassium acetate, 0.1 mg/ml BSA), 2 mM Spermidine, 1 mM non-limiting rNTPs, 20 nM DNA template, 5 mM DTT, 20 µM BI, 0.5 kU/ml RiboLock, 0.25 U/ml PPase, 2 kU/ml, 5 µg/ml NDP kinase, 0.5 kU/ml SP6 RNAPol.

- Master mix for **GMP+GDP+GTP** quantification
  Recipe for 100 reactions (10 µl final reaction volume: 5 µl master mix and 5 µl sample):
  67.5 µl $H_2O$
  100 µl 10x NEB CutSmart buffer
  100 µl of 10 mM non-limiting rNTP mix.
  25 µl of 0.8 µM DNA template for SP6 RNA pol.
  50 µl of 100 mM DTT
  25 µl of 80 mM spermidine
  20 µl of 1 mM BI
  12.5 µl of 40 kU/ml RiboLock
  25 µl of 10 U/ml PPase
  25 µl of 0.2 mg/ml NDP kinase (BioNukleo, NDPK-33)
  25 µl of 0.2 mg/ml GMP kinase (Bionukleo, NMPK-21)
  25 µl of 20kU/ml SP6 RNA polymerase

Final concentrations (after addition of sample): 1x reaction buffer (20 mM Tris-acetate pH 7.9, 10 mM Mg-acetate, 50 mM potassium acetate, 0.1 mg/ml BSA), 2 mM Spermidine, 1 mM non-limiting rNTPs, 20 nM DNA template, 5 mM DTT, 20 µM BI, 0.5 kU/ml RiboLock, 0.25 U/ml PPase, 2 kU/ml, 5 µg/ml GMP kinase, 5 µg/ml NDP kinase, 0.5 kU/ml SP6 RNAPol.

#### Measurement procedure

Set up the reactions on ice. We recommend pipetting of both the sample and the master mix in reverse mode for the maximum precision. Duplicate or triplicate measurements are generally not required for precise quantification of rNTP with a good pipetting technique. Triplicate measurements are, however, recommended when assessing rNMPs and rNDPs due to much lower concentrations of these nucleotides in many sample types.

1. Pipette 5 µl of sample to the bottom of the well of a qPCR plate.
2. Pipette 5 µl of master mix to the side of the well.
3. Gently tap the plate few times against a table, seal the plate, and spin down [Note2].
   Keep the plate on ice.
4. Program a plate reader [Note3 and Note4] to shake the plate and, record fluorescence upon 470 nm excitation and 510 nm emission at 37°C: first 10 min with the minimum possible interval and then 5-8h with the interval of 10 min. The run can be stopped when the reactions have reached the plateau phase (typically 2-5 h).
5. Determine the nucleotide concentrations from baseline-subtracted end-point fluorescence values [Note4].

### Notes

Note 1:
The optimal basal concentration of limiting nucleotide may require slight adjustment depending on the source of the rNTPs.

Note2:
Typical qPCR plate seals have a pressure-sensitive adhesive. If these types of sealing membrane are not carefully applied with sufficient pressure, edge artefacts may take place due to evaporation.

Note3:
Instead of plate reader, a typical qPCR instrument can also be used (SYBR green/FAM channel). qPCR instruments provide better temperature control but somewhat inferior fluorometry in comparison to a dedicated modern plate reader. We usually observe slightly less variation with a plate reader (Biotek Synergy H1) than with a qPCR instrument (Bio-Rad CFX384).

Note4:
The baseline should be determined typically within 2 to 7 minutes of moving the plate to +37°C. The basal fluorescence of BI is somewhat sensitive to temperature, and therefore a few minutes is required for stabilization of the baseline fluorescence. If a temperature gradient is observable regarding the baseline, the plate can be preheated in a PCR block.
