## Supplementary material for "Quantification of all 12 canonical ribonucleotides by real-time fluorogenic *in vitro* transcription": Step-by-step protocol - sample extraction

### Extraction of nucleotides and total protein fraction

(Purhonen and Kallijärvi 2023)

#### Reagents

- Ice-cold 80% MeOH:
- Ice-cold 100% MeOH
- Chloroform. Volatile and toxic, use fume hood! Do not use serological pipettes made of polystyrene to dispense chloroform!
- Diethyl ether (required only for option 2 in the protocol). Volatile and toxic, use fume hood! Do not use serological pipettes made of polystyrene to dispense diethyl ether!
- Milli-Q-grade water
- 1.5 ml microtube
- 2 ml microtubes
- Microtube homogenizer for soft tissues (e.g. Nippon Genetics cat. no. NG010).
- Tissue grinder for fibrous tissues (e.g. WHEATON® Tenbroeck Tissue Grinder, WVR cat. no. 62400-493).
- Microtip probe sonicator.
- SDS buffer: 2% SDS, 1% β-mercaptoethanol, 12% glycerol, and 60 mM Tris-Cl pH 6.8.

#### Extraction of polar metabolites from snap-frozen tissue

1. Cut 10-25 mg of frozen tissue into a 1.5 ml microtube. Keep the samples dry ice cold.

2. Homogenize soft tissues (e.g. liver and kidney) in 0.5 ml of ice-cold 80% MeOH using a microtube pestle homogenizer. Place the homogenized sample back on dry ice before proceeding to the next sample.

   For fibrous tissues (e.g. skeletal muscle), we recommend a more harsh homogenization method e.g. roughened glass-to-glass potter homogenizer.

3. Sonicate the homogenate for 12s with a probe sonicator and appropriate amplitude/power setting for the volume (12% amplitude with Branson Digital Sonifier 250). Place the sonicated sample back on dry ice.

4. Incubate 3 min 100°C in a heat block [Note 1 and 2].
   Place the tubes into the heat block gaps open. Close the gaps after 30-60s.
   Use only microtubes with tight-fitting gaps!

5. Place the sample on wet ice.
   Add 200 µl chloroform. Briefly vortex.

6. Add 200 µl $H_2O$.
   Vortex 10s full speed. Shake vigorously for 30s.

7. Centrifuge 3 min 18000g at 0-4°C.

8. [Optional] Discard most of the lower phase (chloroform-MeOH) with a syringe (same syringe can be used for all samples). Centrifuge 3 min 18000g at 0-4°C. This step facilitates the collection of the upper phase (MeOH-$H_2O$) in the next step.

9. Collect the upper phase (MeOH-$H_2O$) into a new tube.
   Use pre-weighted 2 ml microtubes if continuing with option 2 in the next step.

10. [Optional] Save the interphase and the lower phase for collection of proteins.

11. [Optional] If any protein precipitate is carried into the aqueous phase, clarify the solution by centrifugation.

Option 1

Evaporate the extracts to dryness in a centrifugal vacuum evaporator [Note 3].

Option 2

Wash the aqueous fractions with diethyl ether to remove residual MeOH.

a) Add 1.2 ml of diethyl ether.
   Vortex for ~5s.
   Briefly centrifuge (e.g. 1s 14000g).
   Discard most of the upper layer.

b) Repeat step a) three times. Approximately 300 µl of aqueous extract should be remaining.

c) Evaporate any residual diethyl ether layer under $N_2$ gas flow for a few seconds. If a source of $N_2$ gas is not available, keep the sample tubes gaps open at room temperature until the remaining diethyl ether layer has evaporated (a few minutes).

d) Evaporate the remaining traces of diethyl ether in a centrifugal vacuum evaporator for 6-20 min. If a centrifugal vacuum evaporator is not available, place the sample tubes, gaps open, into a heat block preheated to 90°C for 2 min.

e) Weigh the remaining amount of liquid (1 mg ~ 1 µl) to estimate the volume of the extract.

12. Store the extracts at -80°C.

#### Extraction of polar metabolites from cultured cells

a. Wash the cells once with ice-cold PBS.
b. Scrape the cells directly in an appropriate volume of ice-cold 80% MeOH (0.5 ml per 6 cm dish)
c. Continue from step 3 of the extraction protocol for tissue samples. If necessary, adjust the volumes of the solvents according to the initial volume of 80% MeOH.

#### Collection of total protein fraction for normalization

1. Add 800 µl of ice-cold 100% MeOH to the chloroform-interphase fraction from step 10. Mix.

2. Centrifuge at least 3 min 18000g at 0-4°C
   Remove the supernatant.

3. Centrifuge 1s 18000g.
   Remove any remaining liquid.

4. Add an appropriate volume of SDS buffer (typically 500 µl for 20 mg tissue samples or 150 µl for 6 cm cell culture dishes).

5. Sonicate to completely dissolve the proteins.

6. Incubate 5 min at 95°C.

7. [Optional, sample type-specific step] Clarify the proteins solutions by centrifugation if clear insoluble material persists.

8. Measure protein concentration with a method that is compatible with the SDS buffer [Note4]. We recommend a modified SDS-compatible Bradford assay (Rabilloud 2018, doi.org/10.1371/journal.pone.0195755).

#### Notes

Note 1:
The airspace of the microtubes should be at room temperature or warmer before closing the gaps. Heating sealed initially dry ice-cold tubes will cause the gaps to pop up. If microtube gaps still stubbornly pop up in the heat block, it is possible to pinch a hole into the gap with a 27g syringe. This small hole does not compromise vortexing.

Note 2:
If the total protein fractions are to be used for Western blot or other protein analyses, it is possible to perform the heat denaturation step after step 9 (MeOH-$H_2O$ phase after phase separation) followed by centrifugation. However, this modification may slightly affect the extraction efficiency of protein-bound nucleotides.

Note 3:
Aqueous methanolic extracts are prone to bumping (boiling over/foaming) during vacuum evaporation. This is somewhat a random process and affected by the extract composition and specifications of the evaporator. If a vacuum controller is available, we recommend the following settings: gradient of 250 mbar to maximum vacuum in 40 min, followed by maximum vacuum until

dry. Another option to prevent accidental sample losses during evaporation is to close the tubes with perforated gaps (e.g. a syringe-pinched hole).

[Note4] Glycerol and β-mercaptoethanol can be removed form the buffer if they interfere with the chosen protein assay (e.g.BCA assay).
