## Supplementary methods figures and tables for "Quantification of all 12 canonical ribonucleotides by real-time fluorogenic *in vitro* transcription"

### Extraction of nucleotides for data in Figure 4A, and Supplementary Figures S3 and S4

Frozen liver samples (18-20 mg) were directly homogenized in 0.5 ml of ice-cold 60% MeOH with a battery-operated microtube pestle. The homogenization was finalized with a microtip probe sonicator (12s, amplitude 10%, Branson Digital Sonifier 250). The samples were kept on dry ice between the homogenization steps. Per 0.5 ml of 60% MeOH, 225 µl of chloroform was added to further precipitate macromolecules and to remove non-polar metabolites. The extracts were vortexed for 10s followed by vigorous shaking for 30s. Phase separation was induced by centrifugation (3 min 180000g at 0°C). The aqueous phase was transferred into a new microtube. MeOH was removed by washing three times with 1.4 ml of diethyl ether. Any remaining layer of diethyl ether was evaporated by 10s flow of $N_2$ gas. The traces of the diethyl ether from the aqueous phase were evaporated at 65°C for 5 min in SpeedVac Plus SC110A (Figure 4A and Supplementary Figure S4) or at 25-30°C in Genevac miVac Duo (Supplementary Figure S3) vacuum centrifugal evaporators. The remaining liquid volume after evaporation was estimated by weighing the extracts and assuming the density of 1 mg/µl.

**Supplementrary Table S1**. Reagents and materials

| Reagent | Catalogue number | Source |
|---|---|---|
| T7 RNA polymerase | M0251 | New England BioLabs |
| T7 P266L mutant RNA polymerase (from highYield T7 P&L RNA Synthesis Kit) | RNT-201 | Jena Bioscience |
| SP6 RNA polymerase | M0207 | New England BioLabs |
| ATP, UTP, CTP, GTP | RNT-201 or N0450 | Jena Bioscience or New England BioLabs |
| CutSmart buffer | B6004S or B7204 | New England BioLabs |
| Spermidine | S2626 | Sigma-Aldrich |
| Dithiothreitol (DTT) | RNT-201 or D0632 | Jena Bioscience Sigma-Aldrich |
| BI | 7466 | Tocris Bioscience |
| AMP | 1025 | Jena Bioscience |
| UMP | 1033 | Jena Bioscience |
| CMP | 1032 | Jena Bioscience |
| GMP | 1028 | Jena Bioscience |
| ADP | A5285 | Sigma-Aldrich |
| UDP | 3111 | Tocris Bioscience |
| CDP | C9755 | Sigma-Aldrich |
| GDP | G7127 | Sigma-Aldrich |
| RiboLock | EO0382 | Thermo Scientific |
| Pyrophosphatase | EF0221 | Thermo Scientific |
| Nucleoside diphosphate kinase | NDPK-33 | BioNukleo |
| AMP kinase | NMPK-01 | BioNukleo |
| GMP kinase | NMPK-21 | BioNukleo |
| UMP-CMP kinase | NMPK-22 | BioNukleo |
| 384-well qPCR plate | HSP3805 | BioNukleo |
| Microseal 'B' PCR Plate Sealing Film | MSB1001 | BioNukleo |

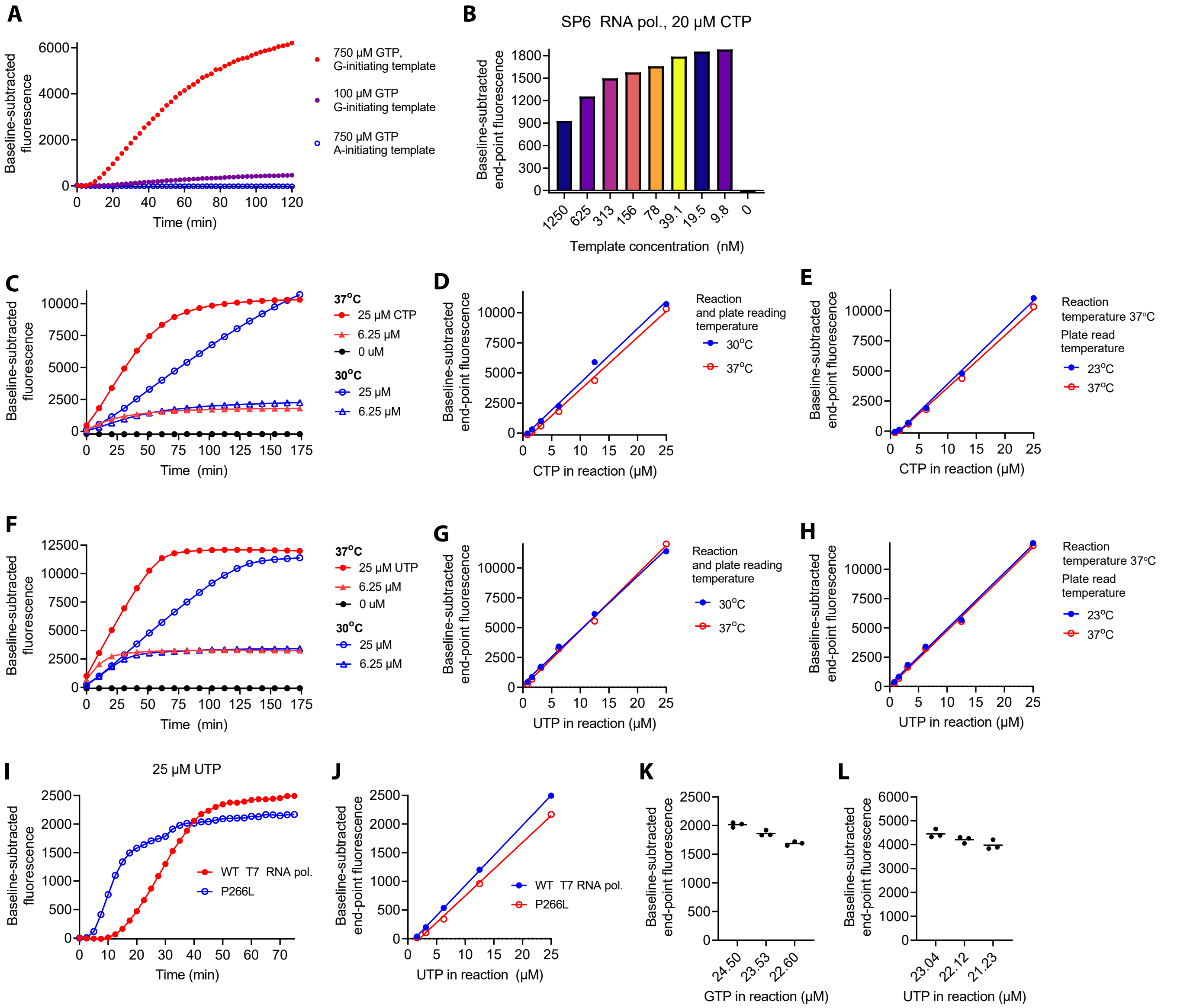


**M**

| Graph | RNA polymerase | Template concentration (nM) | BI (µM) | Reaction temperature (°C) | Reaction volume (µl) | Instrument |
|---|---|---|---|---|---|---|
| A | *P266L T7 | 125 | 10 | 30 | 4 | Plate reader |
| B | 2 kU/ml SP6 | 0 to 1250 | 20 | 37 | 4 | Plate reader |
| C-E | 0.5 kU/ml SP6 | 20 | 20 | 30 or 37 | 4 | qPCR instrument |
| F-H | 1.25 kU/ml T7 | 20 | 20 | 30 or 37 | 4 | qPCR instrument |
| I-J | 5 kU/ml WT T7<br>* P266L T7 | 125 | 10 | 30 | 4 | Plate reader |
| K | 0.5 kU/ml SP6 | 20 | 20 | 37 | 8 | Plate reader |
| L | 1.25 kU/ml T7 | 20 | 20 | 37 | 8 | Plate reader |

Constant variables:
1 mM non-limiting rNTPs, 5 mM DTT, 2 mM spermidine, 25 U/ml PPase, 1x CutSmart buffer
* The P266L T7 RNA polymerase was diluted into the reactions as recommended by the manufacturer (specific activity unkown).

**Supplementary Figure S1.** Optimization of the assay. (**A**) Suitability of A-initiating template for GTP quantification with T7 RNA polymerase. The A-initiating template construct contained the following class II T7 RNA polymerase promoter: 5′-TAATAC-GACTCACTATT-3′. The pre-aptamer leader sequence was 5′-AAATACAAAA-3′ The G-initiating template is described in Table 1 (Template 1). (**B**) Titration of the template concentration for quantifications with SP6 RNA polymerase. (**C**-**E**) Effect of reaction temperature and temperature at which the end-point fluorescence was read on assay sensitivity with SP6 RNA polymerase: (**C**) Real-time monitoring of CTP quantification reactions at 30 and 37°C, and the (**D**) corresponding standard curves from end-point fluorescence values; (**E**) Standard curves from reactions performed at 37°C and the end-point fluorescence read at 23 or 37°C. (**F**-**H**) The experiment in **C**-**E** repeated with T7 RNA polymerase. (**I**-**J**) Comparison of wildtype (WT) and P266L mutant T7 RNA polymerase for quantification of UTP. (**K**-**L**) Testing of assay resolution by quantification of approximately 4% differences in limiting rNTP concentration. (**M**) Description of the assay conditions.

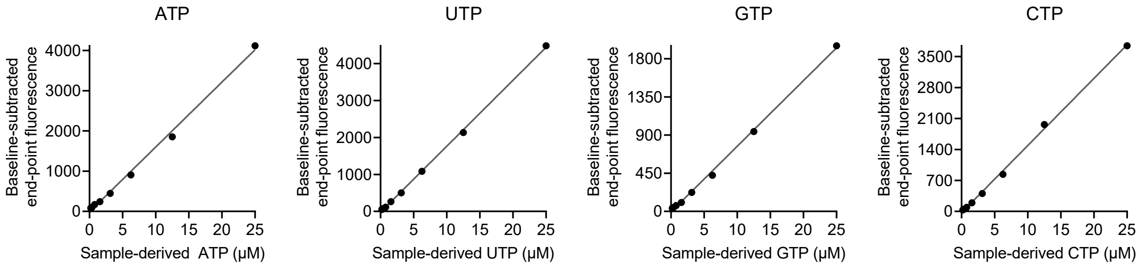


**Supplementary Figure S2.** Low basal concentrations of limiting nucleotide linearize the standard curves with monomeric Broccoli-based detection. The basal limiting nucleotide concentrations were 1 µM for quantification of ATP, UTP, CTP, and 2 µM for quantification of GTP.

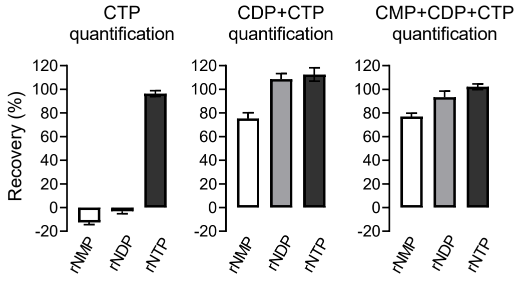


**Supplementary Figure S3.** Interfering rNMP kinase activity in unboiled liver extracts. Liver extracts were spiked with rNMP, rNDP, or rNTP mixtures comprising: 2000 pmol adenosine nucleotides (AMP, ADP, or AMT), 200 pmol uridine nucleotides (UMP, UDP, or UTP), 200 pmol guanosine nucleotides (GMP, GDP, or GTP), and 50 pmol cytidine nucleotides (CMP, CDP, or CTP) per mg tissue. Signal increase due to exogenous CMP, CDP, and CTP was assessed in CTP, CTP+CDP, and CTP+CDP+CMP quantification reactions. Diethyl ether washed extracts with a short vacuum evaporation step at room temperature were used for these experiments (Supplementary methods). The bar graphs represent mean and SEM from 4 independent liver extracts.

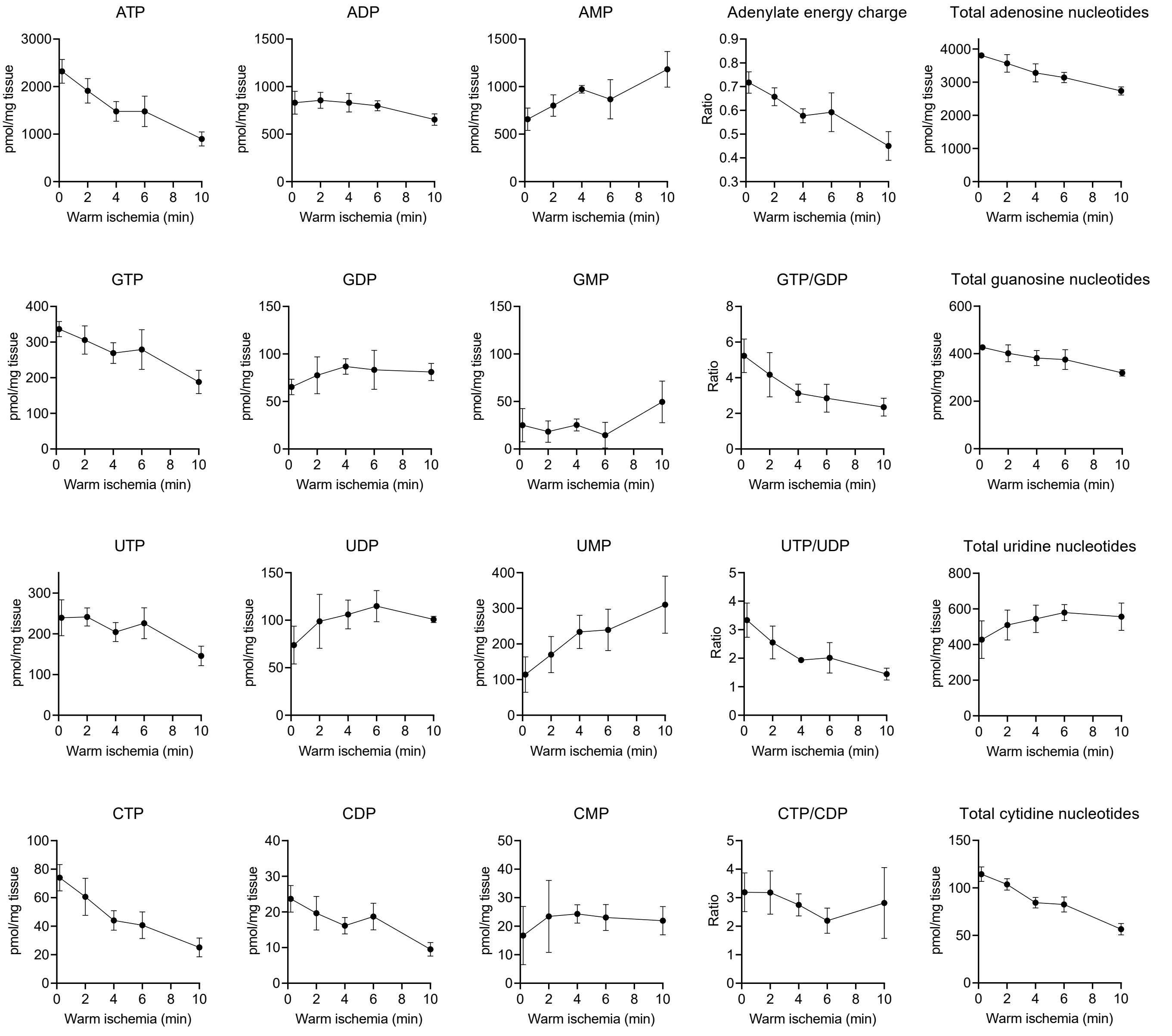


**Supplementary Figure S4.** Warm ischemia experiment related to Figure 5. For this experiment, the liver extracts were prepared as described in Supplementary Methods. In brief, the extracts were exposed for 5 min to 65°C in vacuum centrifugal evaporator but did not undergo controlled heat denaturation step as described in Materials and Methods. These data are potentially affected by residual sample-derived enzymatic activity. The data points represent mean and SD from four non-fasted male mice of 2 month of age.
